# Caloric Restriction recovers impaired β-cell-β-cell coupling, calcium oscillation coordination and insulin secretion in prediabetic mice

**DOI:** 10.1101/2020.03.03.975961

**Authors:** Maria Esméria Corezola do Amaral, Vira Kravets, JaeAnn M. Dwulet, Nikki L. Farnsworth, Robert Piscopio, Wolfgang E. Schleicher, Jose Guadalupe Miranda, Richard K. P. Benninger

## Abstract

Caloric restriction has been shown to decrease the incidence of metabolic diseases such as obesity and type 2 diabetes mellitus (T2DM). The mechanisms underlying the benefits of caloric restriction involved in insulin secretion and glucose homeostasis and are not fully understood. Intercellular communication within the islets of Langerhans, mediated by Connexin36 (Cx36) gap junctions, regulates insulin secretion dynamics and glucose homeostasis. The goal of this study was to determine if caloric restriction can protect against decreases in Cx36 gap junction coupling and altered islet function induced in models of obesity and prediabetes. C57BL6 mice were fed with a high fat diet (HFD), showing indications of prediabetes after 2 months, including weight gain, insulin resistance, and elevated fasting glucose and insulin levels. Subsequently, mice were submitted to one month of 40% caloric restriction (2g/day of HFD). Mice under 40% caloric restriction showed reversal in weight gain and recovered insulin sensitivity, fasting glucose and insulin levels. In islets of mice fed the HFD, caloric restriction protected against obesity-induced decreases in gap junction coupling and preserved glucose-stimulated calcium signaling, including Ca^2+^ oscillation coordination and oscillation amplitude. Caloric restriction also promoted a slight increase in glucose metabolism, as measured by increased NAD(P)H autofluorescence, as well as recovering glucose-stimulated insulin secretion. We conclude that declines in Cx36 gap junction coupling that occur in obesity can be completely recovered by caloric restriction and obesity reversal, improving Ca^2+^ dynamics and insulin secretion regulation. This suggests a critical role for caloric restriction in the context of obesity to prevent islet dysfunction.

## Introduction

Type 2 diabetes mellitus (T2DM) results from environmental factors such as diet and absent physical activity that contribute to insulin resistance, as well as a genetic predisposition that reduces islet compensation (Byrd et al., 2018; Zheng et al., 2018). There is a strong link between obesity, weight gain and T2DM (Franks and McCarthy, 2016; Mansourian et al., 2018). Diets high in fat have been indicated to be contributory factors in the development and progression of T2DM. For example, excess exposure of pancreatic islets to lipids (lipotoxicity) contribute to the loss of β-cell mass and function (Ježek et al, 2018; Imai et al., 2019). It is possible that the excess of lipids in the diet result in higher cholesterol, triglycerides and free-fatty acid levels that may lead to impaired β-cell secretory function, metabolism, and viability (Bugliani et al, 2019).

In the islet, gap junctions composed of Connexin36 (Cx36), provide electrical coupling between β-cells that promote regulation of electrical activity and insulin secretion (Calabrese et al., 2004; Head et al, 2012; Benninger et al., 2011; Ravier et al., 2005; Speier et al. 2007). A key role for gap junctions are to allow for synchronous free-calcium ([Ca^2+^]_i_) oscillations across the islet, which is driven by the electrical coupling between adjacent β-cells (Calabrese et al., 2003; Benninger et al 2008; Westacott et al, 2017). In Cx36 deficient mice, synchronized [Ca^2+^]_i_ oscillations are no longer observed at the whole-islet level (Ravier et al., 2005; Benninger et al., 2008). These mice also show disrupted first phase and pulsatile second phase insulin release and are glucose intolerant (Head et al., 2012). These changes in insulin secretion observed in Cx36 deficient mice are similar to those observed in prediabetic and type 2 diabetic states, indicating that Cx36-dependent signaling may be disrupted under pre-diabetic conditions.

Lipotoxicity has been indicated to disrupt Cx36 gap junction within the islet under ex-vivo based conditions (Allagnat et al. 2008; Allagnat et al, 2005; Carvalho et al., 2012, Haefliger et al, 2013). Similarly, synchronous [Ca^2+^]_i_ oscillations are disrupted under exvivo lipotoxic conditions and in islets from ob/ob mice (Ravier et al 2005; Hodson et al., 2013). As such the disruption to islet function and insulin release dynamics upon a loss of Cx36 gap junction coupling may contribute to diabetes progression. Furthermore, a polymorphism in the *GJD2* gene that encodes Cx36, is associated with altered β-cell function and associated with type 2 diabetes (Cigliola et al.,2016). This further suggests a key role Cx36 gap junctions may play in the link between obesity, islet dysfunction and type2 diabetes.

Obesity can be mediated by diet, but also includes a genetic predisposition and can be modified pharmacologically (Byrd et al., 2018). Given the importance that nutrition exerts in the appearance of metabolic diseases, many studies involving caloric restriction (CR) have been performed, demonstrating that this intervention can modulate pathways preventing metabolic diseases (López-Domènech et al, 2019; Madeo et al., 2019; Lean et al. 2019). Dietary restriction preserved pancreatic β-cell function and β-cell mass in diabetic db/db mice, suppressing cellular apoptosis and antioxidative stress in the pancreatic β cells (Yan et al, 2018; Kanda et al., 2015). More broadly, ovariectomized rats submitted to short term CR of 40% have shown enhanced muscarinic response (which promotes insulin release) and antioxidant defenses (Pacher et al., 2019). In contrast lean rats submitted to CR of 40% show little change in intracellular [Ca^2+^]_i_ concentration in isolated pancreatic islets (do Amaral et al., 2011).

Given the enhancement of islet function in obesity or diabetes under CR, the goal of this work was to investigate whether CR promotes alterations in Cx36 gap junction coupling and associated [Ca^2+^]_i_ and insulin secretion regulation in the islet during the altered function that occurs under obesity and lipotoxic conditions. Using a model of diet-induced obesity, and mice submitted to 40% CR, we found that CR overcomes high fat diet induced chnages to recover islet gap junction coupling, [Ca^2+^]_i_ dynamics and insulin secretion; thus showing how short term CR is sufficient to recover islet function.

## Materials and Methods

### Ethical Approval

All animal procedures and experiments were performed in accordance with protocols approved by the Institutional Animal Care and Use Committee at the University of Colorado Anschutz Medical Campus (B-95817(05)1D).

### Experimental Animals

Male C57/BL6 mice were purchased from Jackson Laboratories (Bar Harbor, ME). Animals were housed at 22±1°C on a 12:12-h light-dark cycle and had free access to water. At age 8 weeks, the animals were divided into three groups. One group was fed 12 weeks with a standard rodent diet (ENVIGO 2020X) containing 16% lipids, 60% carbohydrates, and 24 % proteins with 3.1 Kcal/g (normal diet, ND). The second group was fed for 12 weeks with high-fat-diet (ENVIGO TD.03584) containing 35.2% lipids (approximate fatty acid profile: 40% saturated, 50% monounsaturated, 10% polyunsaturated), 36.1% carbohydrates, and 20.4 % proteins with 5.4Kcal/g (high-fat diet, HFD). The third group was fed with HFD for 8 weeks, and then for 4 weeks received a restricted amount of the HFD diet to achieve caloric restriction (HFD-CR). This restriction was achieved by providing animals with 60% of the daily consumption of the HFD group, on a daily basis. This corresponded to a moderate restriction of 40% of the control intake (2 g/day) (Stankovic et al 2013).

### Glucose and insulin tolerance tests (IPGTT, ITT)

The intra-peritoneal glucose tolerance test (IPGTT) was performed after fasting the animals for 6 h (n=6 animals per group) at 11 weeks during the experiment (Ayala et al, 2010). Glucose was administered to the animals intraperitoneally (2g/kg body weight in sterile PBS). Blood samples were collected from the tail, before glucose administration (time zero) and at 15, 30, 60, 90, and 120 min. after glucose administration. The blood glucose was determined using a glucose meter (Contour® Ascensia, Beyer). The insulin tolerance test (ITT) was performed after fasting the animals for 6 h. Human recombinant insulin (Apidra® insulin glulisine, Sanofi Aventis US LLC, Bridgewater, NJ) was administered to the animals intraperitoneally (1.5 U/kg body weight). Blood glucose was sampled as above at time zero (before insulin injection) and at times of 5, 10, 15, 20, 25, and 30 min. The glucose disappearance constant (K_itt_) was calculated using the formula: ln(2)/t_1/2_. The serum glucose t_1/2_ value was calculated from the slope of a linear regression applied over the linear phase of the glucose decrease (0-15min) (Bonora et al., 1989).

### Plasma insulin measurements

Blood (~50μl) was collected after fasting the animals for 6 h. Blood was centrifuged to obtain serum, and insulin concentration determined by ELISA (Mouse ultrasensitive insulin ELISA, #80-INSMSU-E10; Alpco, Salem NH), following the manufacturer’s instructions.

### Islet Isolation and Culture

Pancreata were isolated from mice after 12 weeks of the study. Mice were first anesthetized by intra-peritoneal injection of 80mg/kg ketamine (Zetamine™, NDC 13985-702-10; Vet One / MWI, Boise, ID) and 20mg/kg xylazine (AnaSed LA, NDC 13985-704-10; Vet One / MWI), and euthanized by exsanguination. Collagenase (type V, Sigma Aldrich) was injected into the pancreas through the pancreatic duct, the pancreas was isolated immediately following infusion, the pancreas incubated at 37°C and islets hand-picked. Islets were incubated overnight in 1640 RPMI Medium (Sigma) with 10% FBS, 10,000 U/mL Penicillin and 10,000 μg/mL Streptomycin at 37°C with 5% CO_2_.

### Fluorescent Recovery After Photobleaching (FRAP)

Islets were cultured on MatTek dishes (MatTek Corp., Ashland, MA) coated with CellTak (BD Biosciences, San Jose, CA) overnight. Prior to imaging, islets were stained with 12.5μM rhodamine 123 (R123, Sigma Aldrich) for 30 min at 37degC, in imaging buffer containing 125mM NaCl, 5.7mM KCl, 2.5mM CaCl2, 1.2mM MgCl2, 10mM HEPES, 0.1% bovine serum albumin (BSA), and 2mM glucose. Islets were imaged at room temperature on a Zeiss LSM 800 confocal microscope with a 40x 1.0 NA water immersion objective. R123 was excited or photobleached with a 488nm laser. FRAP was performed as previously described (Farnsworth et al., 2014). Briefly, half of each islet was photobleached for 360s, and the recovery of fluorescence in the bleached area was measured over time. Recovery rates were calculated from the inverse exponential fluorescence recovery curve *I(t)* = *(I_∞_−I_p_) (1−e^−kt^)* + *I_p_* for the entire bleached area.

### Intracellular [Ca^2+^]_i_ Imaging and Analysis

Islets were cultured on MatTek dishes coated with CellTak overnight. Prior to imaging, islets were stained with 4μM Fluo-4 AM (Thermo Fisher Scientific) in imaging buffer (see above) at 2mM glucose, at room temperature. Islets were imaged at 37°C on a Zeiss 800 LSM confocal microscope with a 40x 1.0 NA water objective. Images were taken every second for 5 minutes with 2mM glucose, and after stimulation with 11mM glucose for 10 minutes. The fraction of the islet which responded to glucose stimulation with oscillatory increases in intracellular [Ca^2+^]_i_ (active area), the fraction of the islet which showed coordination of the oscillatory increases in intracellular [Ca^2+^]_i_ (fraction correlated area), and the oscillation frequency were determined using MatLab (MathWorks, Natick, MA) as previously described (Hraha et al., 2014).

### NAD(P)H analysis

Islets were cultured on MatTek dishes coated with CellTak overnight. Prior to imaging islets were incubated at 2mM glucose in imaging buffer (see above). NAD(P)H autofluorescence was imaged under two-photon excitation using a tunable mode-locked Ti:sapphire laser (Chameleon; Coherent, Santa Clara, CA) set to 720 nm. Fluorescence emission was detected at 400–450 nm using the internal detector. Z-stacks of 10 images were acquired spanning a depth of 10 μm. The average NAD(P)H intensity was calculated over the volume of the islet imaged. For each day of imaging, the NAD(P)H intensity for each islet over each condition was normalized to the mean NAD(P)H intensity measured at 2mM glucose in ND islets.

### Insulin Secretion Analysis

After overnight incubation, islets were incubated in Krebs-Ringer buffer with 0.1% BSA and 2mM glucose for 1 hour at 37°C. Five islet equivalents were then incubated in 500μl Krebs buffer at either 2mM and 20mM glucose for 1 hour. Supernatant was then collected for analysis of insulin secretion. To assess content, islets were lysed by freezing in 2% Triton-X-100 in deionized water. Insulin was measured with a mouse ultrasensitive insulin ELISA kit (Alpco) per the manufacturer’s instructions.

### Islet Viability

After overnight incubation, islets were stained with propidium iodide (Sigma Aldrich) to detects dead cells (displayed in red), and fluorescein diacetate (Sigma Aldrich) to detect live cells (displayed in green). Islets were then imaged immediately on a Zeiss 800 LSM confocal microscope at three depths (every 3μm).

### Statistical analysis

Comparative analysis of the results for the different groups was performed using ANOVA followed by Tukey’s post-test. The results were expressed as mean ± standard error of the mean (±SEM), and the level of significance adopted was 5% (p<0.05). When comparing two groups a t-test was used, as explicitly stated.

## Results

### Caloric Restriction (CR) reverses high fat diet induced obesity and prediabetes

We first established and characterized the models of obesity and caloric restriction (CR). Figure 1A shows the experimental timeline. Mice were put on a diet that was high in fat (high fat diet, HFD) or remained on a normal chow diet (ND). Following 8 weeks on HFD, a subset of the HFD group underwent caloric restriction (CR) for 4 weeks (HFD-CR). After 8 weeks, the HFD feeding induced a significant increase (* p=0.02) in the body weight of mice compared with the ND group (Fig. 1B). HFD-CR promoted a significant reduction in the body weight of mice under the last 2 weeks compared with the HFD group (**p=0.02). The HFD-CR group showed lower food consumption than the ND group and unrestricted HFD group, as expected (p<0.0001; Fig. 1C). Consistent with diet-induced weight gain, HFD mice showed increased peri-epididymal fat pad mass, which was significantly reduced in the HFD-CR group (p=0.0084; Fig. 1D). Fasting blood glucose and fasting plasma insulin levels were both significantly elevated in the HFD group compared to the ND group (p=0.031, Fig. 1E; p=0.0032, Fig.1F). Fasting blood glucose and fasting plasma insulin levels were significantly reduced in the HFD-CR group compared to the HFD group (p=0.002, Fig.1E; p<0.0001, Fig.1F). Thus HFD mice showed features of obesity and pre-diabetes that were reversed following 4 weeks CR.

**Figure 1.**
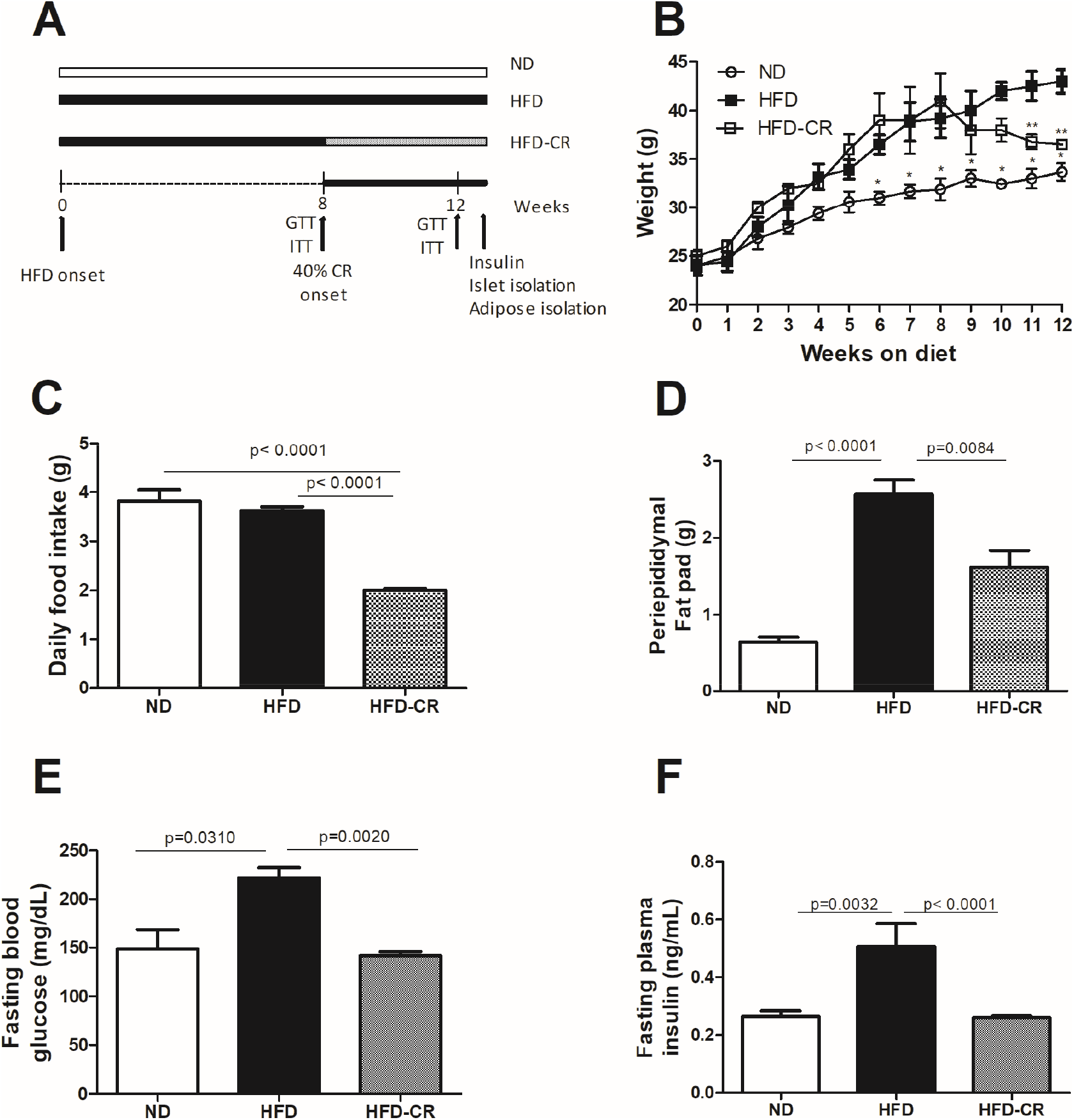
Establishing caloric restriction reversal of obesity. (A) Timeline to show application of diets in 3 mice groups, together with assays used. One group consume a standard rodent diet (normal diet-ND). The second group consume a high-fat-diet (HFD). The third group received HFD and then 60% of the HFD food, corresponding to a moderate caloric restriction of 40% of the HFD (HFD-CR). (B) changes in body weight (g) of the animals. (C) Daily food intake (g). (D) Periepidydimal fat pad (g). (E) fasting glycemia (mg/dL). (F) fasting insulin (ng/mL) for mice of the ND, HFD and HFD-CR groups (n=6 mice per group). Data represent mean ± SEM. *(p<0.05) to ND vs HFD and ** (p<0.05) to HFD vs HFD-CR based on one-way ANOVA with Tukey post-test.

### CR improves glucose tolerance and insulin sensitivity

Given reversal of elevated fasting insulinemia and glycemia upon CR, we next further characterized glucose homeostasis in these mice. After 8 weeks of HFD, prior to CR, mice showed significantly elevated excursions of glucose levels during an IPGTT compared with ND mice (Fig. 2A, B). The HFD and HFD-CR groups showed similar excursions in glucose levels. After 12 weeks of HFD, mice showed an even greater excursion in glucose levels during an IPGTT compared to ND mice (Fig. 2C,D). In contrast following 4 weeks of CR, HFD-CR mice showed a significant decrease in the glucose excursion during an IPGTT compared to HFD mice (Fig. 2C,D), although this was still greater than in ND mice. Thus, CR partially recovers the disruption to glucose tolerance induced by a HFD.

**Figure 2.**
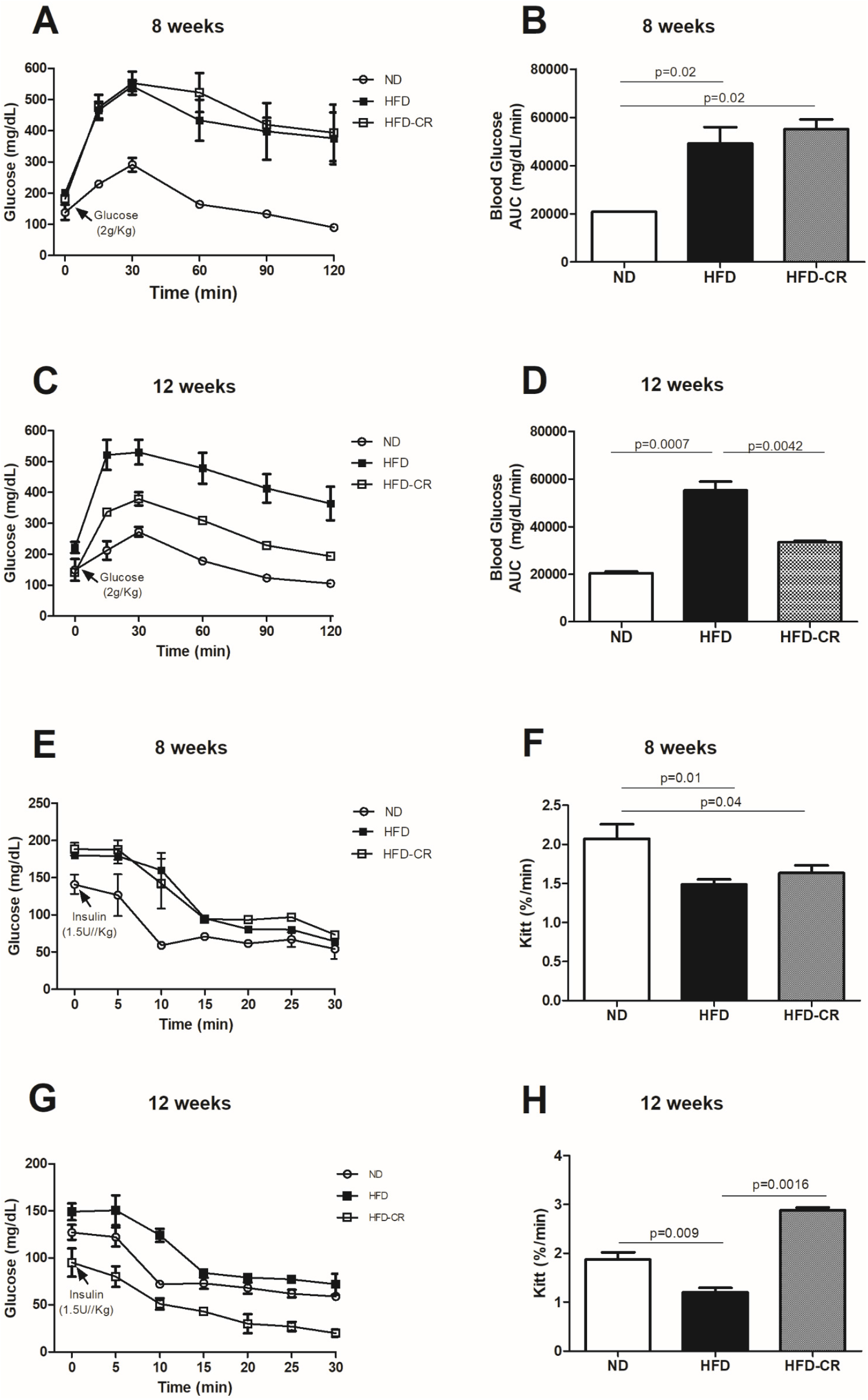
Glucose homeostasis in CR model. (A) Time-course of IPGTT following 8 weeks of high-fat-diet feeding, prior to CR. (B) Area under curve of IPGTT in A. (C) Time-course of IPGTT following 12 weeks of high-fat-diet feeding, after CR imposed. (D) Area under curve of IPGTT in C. (E) Time-course of ITT following 8 weeks of high-fat-diet feeding, prior to CR. (F) Kitt for ITT in E. (G) Time-course of ITT following 12 weeks of high-fat-diet feeding, after CR imposed. (H) Kitt for ITT in G. Data expressed for ND (normal diet), HFD (high fat diet) and HFD-CR (high fat diet caloric restriction) groups (n=6 mice per group). Data represent mean ± SEM, p<0.05 indicates a significant difference based on one-way ANOVA, Tukey post-test.

After 8 weeks of HFD, prior to CR, mice showed significantly slower declines in glucose levels during an ITT compared with ND mice (Fig. 2E,F), indicating reduced insulin sensitivity. The HFD and HFD-CR groups showed similar decreases in glucose levels. After 12 weeks of HFD, mice showed an even slower decrease in glucose levels during an ITT compared to ND mice (Fig. 2G,H). In contrast following 4 weeks of CR, HFD-CR mice showed a significant increase in the rate of glucose lowering during an ITT compared to HFD mice (Fig. 2G,H), which was greater than that observed in ND mice. Thus, CR fully recovers the disruption to insulin sensitivity induced by a HFD.

### CR protects against HFD-induced disruption to islet insulin release

To understand how the above metabolic feature relate to islet function, insulin secretion was first analyzed. At non-stimulatory glucose concentrations (2mM), insulin secretion was similar between all 3 mice groups (Fig. 3A). Islets from HFD mice secreted less insulin at 11mM glucose than ND (p=0.037, Fig. 3A) and HFD-CR islets (p=0.0115, Fig. 3A). As a result, HFD-CR islets in the presence of 11mM glucose had significantly greater glucose-induced insulin secretion than HFD islets (Fig. 3B), which was similar to the glucose-induced insulin secretion in ND islets. We next measured glucose-induced NAD(P)H increases to assess glucose metabolism in each group (Fig. 3C). A slight increase in glucose-induced NAD(P)H autofluorescence was measured HFD-CR islets compared with HFD islets (t-test, p=0.04), consistent with measurements of insulin secretion. However, the decrease in glucose-induced NAD(P)H in HFD islets compared to ND islets was not statistically significant. In contrast to the substantially altered islet function induced by HFD and following CR, we observed no change in islet cell viability (Fig. 3E, F) across all three groups. Thus, CR protects against islet dysfunction induced by HFD.

**Figure 3.**
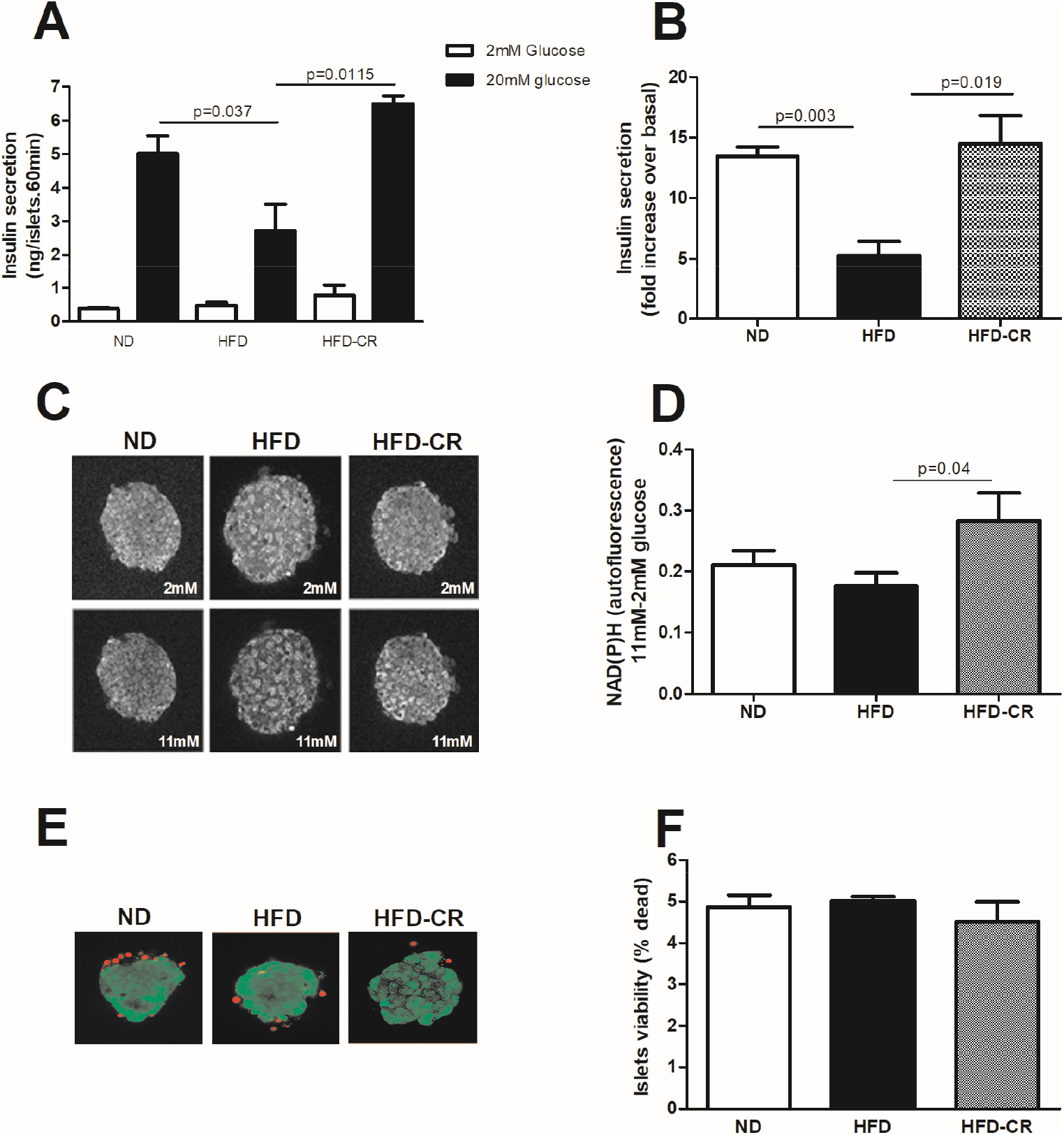
Islet function in CR model. (A) insulin secretion from isolated islets under 2mM (white) and 20mM glucose (black). (B) Fold-change in insulin secretion from 2mM to 20mM glucose, for data in A. (C) Representative images of NAD(P)H autofluorescence at 2mM glucose and after 10min in 11mM glucose. (D) Change in mean NAD(P)H autofluorescence between 2mM and 11mM glucose. (E) Representative images indicating islet viability (red=dead, green=live). (F) Islet viability indicated by % dead cells. Data expressed for islets from ND (normal diet), HFD (high fat diet) and HFD-CR (high fat diet caloric restriction). Data represent the mean ± SEM, p<0.05 indicates a significant difference based on one-way ANOVA, Tukey post-test or Student’s t-test in D.

### CR protects against high fat diet induced decreases in Cx36 Gap Junction

Gap junction coupling is critically important to coordinate β-cell function across the islet, and is disrupted under conditions associated with diabetes such as lipotoxicity (Hodson et al., 2013, Allagnat et al., 2008). We measured Cx36 gap junction permeability by Fluorescence Recovery After Photobleaching (Farnsworth et al 2014). Representative images of the Rh123 fluorescence tracer before photobleaching (0s), immediately after photobleaching (45s), and after fluorescence recovery (360s) are showed (Fig. 4A) Representative fluorescence recovery time courses for each group are also shown (Fig. 4B). Consistent with previous reports of reduced Cx36 protein levels (Carvalho et al., 2012), HFD significantly decreased Cx36 gap junction permeability in HFD islets compared to ND islets, as measured by the rate of Rh123 transfer and fluorescence recovery (p=0.0042; Fig. 4C). HFD-CR islets showed a significant increase in gap junction permeability compared to HFD islets (Fig.4B; p= 0.0205; Fig. 4C). The permeability in HFD-CR islets was not significantly different than in ND islets. Thus, CR recovers the decline in islet Cx36 gap junction permeability induced by HFD.

**Figure 4.**
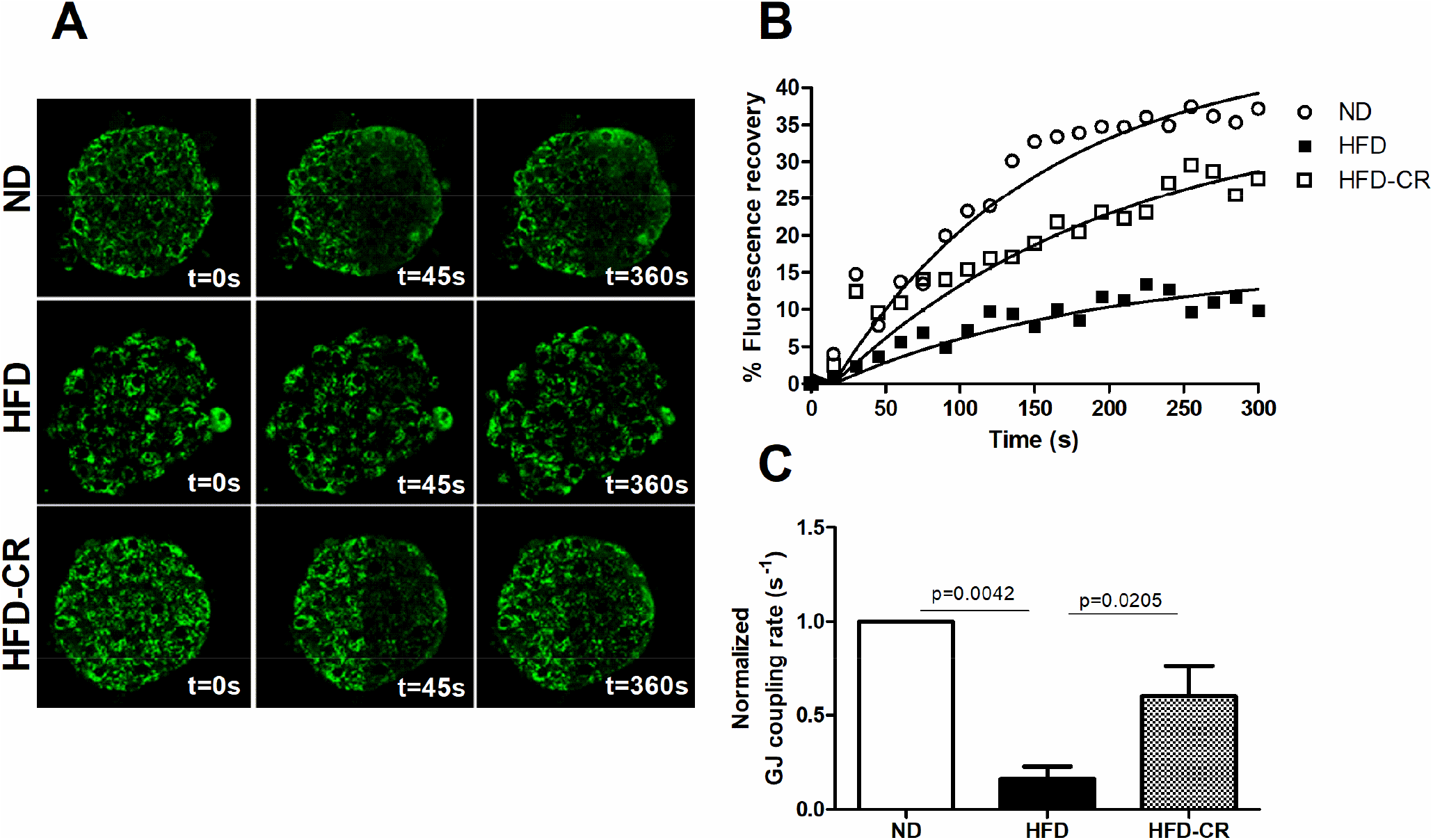
Islet gap junction permeability in CR model. (A) Representative images of Rh123 fluorescence before photobleaching (0s), immediately after photobleaching (45s), and after fluorescence recovery (360s). (B) Representative fluorescence recovery time course for an islet from each experimental group, together with exponential fit. (C) Mean gap junction permeability as expressed by the rate of Rh123 fluorescence recovery. Data expressed for islets from ND (normal diet), HFD (high fat diet) and HFD-CR (high fat diet caloric restriction). Data represent the mean ± SEM over 8-10 islets (from n=6 mice). Data in C is normalized to the mean rate measured in ND islets for each mouse. p<0.05 indicates a significant difference based on one-way ANOVA, Tukey post-test.

### CR protects against high fat diet induced disruptions to islet [Ca^2+^]_i_ oscillations

Gap junction-mediated electrical coupling is required to coordinate intracellular Ca^2+^ and insulin secretion dynamics in the islet (Benninger et al., 2008; Ravier et al. 2005). We next determined if HFD impaired islet Ca^2+^ dynamics, and if CR could recover HFD-induced impairment of [Ca^2+^]_i_ dynamics. Representatives time-courses of Ca^2+^ changes are shown for ND, HFD and HFD-CR islets (Fig. 5A-C). Consistent with previously reports (Carvalho et al., 2012) HFD-feeding increased [Ca^2+^]_i_ oscillation frequency (p=0.0313; Fig. 5D) and decreased [Ca^2+^]_i_ oscillation amplitude (p=0.00195; Fig. 5E) compared to ND islets. Consistent with reduced Cx36 gap junction permeability, HFD did not impact the proportion of cells showing elevated Ca^2+^ oscillations (Fig. 5F) but decreased the proportion of the islet showing coordination of [Ca^2+^]_i_ oscillations (p=0.0013; Fig. 5G) compared to ND islets. CR reversed all effects of HFD-fed changes to [Ca^2+^]_i_ oscillations, including decreasing oscillation frequency (p=0.0424; Fig. 5D), increasing [Ca^2+^]_i_ oscillation amplitude (p=0.0026; Fig. 5E), and increasing the proportion of the islet showing coordination of [Ca^2+^]_i_ oscillations (p=0.0086; Fig. 5G). Thus CR fully reverses disruptions to [Ca^2+^]_i_ dynamics induced by HFD, consistent with the recovery to Cx36 gap junction permeability.

**Figure 5.**
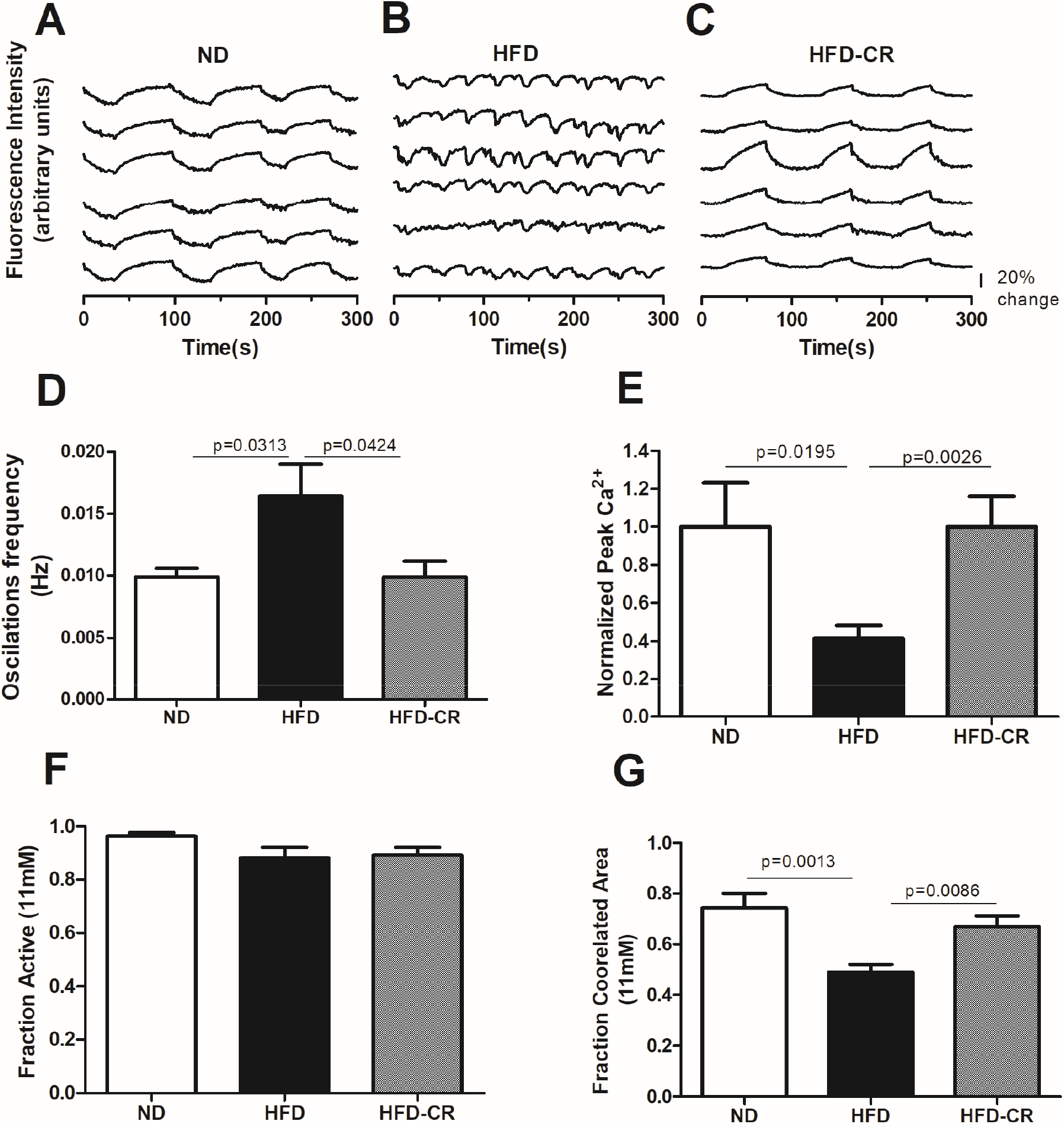
Islet Ca^2+^ in CR model. (A) Representative intracellular [Ca^2+^]_i_ oscillations at 11mM glucose in islets from ND (normal diet). (B) As in A for islets from HFD (high fat diet). (C) As in A for islets from HFD-CR (high fat diet caloric restriction). (D) Ca^2+^ oscillation frequency, (E) Ca^2+^ oscillation amplitude, (F) fraction of islet with elevated Ca^2+^ oscillations at 11mM glucose, (G) fraction of islet with coordinated Ca^2+^ oscillations at 11mM glucose, in isolated islets from ND (normal diet), HFD (high fat diet) and HFD-CR (high fat diet caloric restriction). Data represent the mean ± SEM over 8-10 islets (from n=6 mice) for each condition. Data in E is normalized to the mean amplitude measured in ND islets for each mouse. p<0.05 indicates a significant difference based on one-way ANOVA, Tukey post-test.

## Discussion

Caloric Restriction (CR) has been experimentally performed for the study of physiological and metabolic mechanisms as well as the study of intracellular signaling pathways leading to weight reduction and glucose homeostasis (Golbidi et al, 2017; Veyrat-Durebex et al, 2009). In the islet, physiological insulin secretion is regulated via coordination of membrane depolarization and [Ca^2+^] as a result of Cx36 gap junction channels. Many factors associated with type2 diabetes, including aging (Westacott et al., 2017), lipotoxicity (Allagnat et al., 2008; Hodson et al, 2013), diet (Carvalho et al., 2012), low-grade inflammation (Farnsworth et al. 2016) and genetic predisposition can impact gap junction function and [Ca^2+^] regulation (Cigliola et al. 2016). While dietary restriction has proven effective in enhancing glucose homeostasis in diabetes and pre-diabetes, it is unclear mechanistically how islet function is impacted under such restriction. The aim of this study was to investigate the effects of short-term CR can overcome high fat diet-induced decreases in Cx36 gap junction coupling and altered [Ca^2+^] dynamics. Our key findings were that the coordinated [Ca^2+^] response at elevated glucose is disrupted in pre-diabetic islets, correlating with reduced gap junction coupling and disrupted insulin secretion. However these disruptions were fully recovered by short-term 40% CR.

### Impaired [Ca^2+^] dynamics in HFD mice

As expected, pre-diabetic characteristics were observed in animals fed a high-fat diet (HFD). This included increased body weight and adipose tissue, increased fasting glycemia and insulinemia, but reduced glucose intolerance and decreased insulin sensitivity. During the pre-diabetic state, it is generally understood that compensation is induced within the islet as a result of peripheral insulin resistance. As a result, β-cell mass and insulin output are increased. However, associated with this prediabetic state, are a number of factors that can deleteriously impact islet function, increase ER stress and cause β-cell failure (Swisa et al 2017). In the face of this islet compensation that we observed in the HFD model of prediabetes, a striking feature we observed was the substantial decline in Cx36 gap junction permeability and [Ca^2+^] dynamics.

Declines in Cx36 gap junction protein levels and permeability have been previously reported following similar durations of HFD (Carvalho et al., 2012). This decline in Cx36 gap junctions is similar to the gap junction coupling pattern found in islets submitted to inflammatory cytokines and excess palmitate (Farnsworth et al., 2019, Allagnat et al., 2008). Obesity and cytokines released by adipose tissue are considered a precipitating factor of β-cell dysfunction and also serves as a connection between obesity and T2D (Ciregia et al 2017). Here we also report that islets from HFD showed reduced coordination of [Ca^2+^] oscillations, as well as increased in frequency and reduced amplitude (Fig. 5). Studies in islets from diabetic db/db mice have indicated early disappearance of [Ca^2+^]_i_ oscillations (Gilon et al. 2014). In islets from ob/ob mice smaller amplitude [Ca^2+^] oscillations, with reduced coordination were observed (Ravier et al., 2002; Lee et al. 2018). Pro-inflammatory cytokines, TNF-α and IL-1β are increased in ob/ob and HFD-fed islets (Lee et al. 2018). Islets exposed for 48h to a free fatty acid to model lipotoxicity had reduced [Ca^2+^]_i_ oscillation amplitude (Qureshi et al. 2015). Thus there is a clear link between factors associated with models of obesity and pre-diabetes and the dysregulation of gap junction permeability and Ca^2+^ oscillatory dynamics.

Another indication of islet dysfunction is the impairment of the NAD(P)H elevation that we observed in islets isolated from the HFD mice (Fig. 3C). Declines in NAD(P)H or glucose metabolism in general have been observed in several models of pre-diabetes (Hatanaka et al., 2017), as well as upon other factors such as aging that predispose for type2 diabetes (Gregg et al., 2016). Similar to these results, animals treated with dexamethasone showed an increase in triglyceride, free fatty acids, fasting glycemia and insulinemia also showed reductions in NAD(P)H in isolated islets (Roma et al 2012). The deleterious effects of FFAs have been linked to decreased NAD(P)H content (Iizuka et al, 2002).

As such the HFD mice we examined showed dysfunction to Cx36 gap junction permeability and Ca^2+^ dynamics, as well as other aspects of islet function. This dysfunction is consistent with the effect of a number of factors associated with prediabetes and HFD in prior studies.

### Impact of CR to enhance coordinated [Ca^2+^]_i_ and, gap junction coupling

CR has been performed for the study of physiological and metabolic mechanisms as well as the intracellular signaling pathways leading to weight reduction and improved glucose homeostasis (Golbidi et al, 2017; Veyrat-Durebex et al, 2009). One of the most stable outcomes in animal models of CR is a restitution in glucose tolerance (Matyi et al 2018) and correction of weight by loss of adipose tissue (Dommerholt et al 2018). Adipose tissue exemplifies the most important tissue to adjust for the fast fall in intake (Mitchell et al 2015). In the present study, we confirmed the expected rapid improvement in glucose tolerance with CR, as seen in the HFD-CR group. This included improved fasting glycemia and insulinemia, improved glucose tolerance, improved insulin sensitivity and improved insulin secretion This is consistent with previous work that showed that CR animals presented lower fasting insulinemia (do Amaral et al., 2011; Pires et al., 2014; Mitchell et al 2015; Pacher et al. 2019). These studies utilized alternative models including aging, Ins2^−/−^ and ovariectomized animal models submitted to 40% RC. Therefore, here we can extend these findings to the HFD mouse model

We describe for the first time that CR promotes recovery in factors that are observed to change very early in the progression of diabetes, including the functioning of Cx36 gap junction channels and associated oscillatory [Ca^2+^]_i_ dynamics. This change also correlated with enhanced insulin secretion. Importantly, the complete recovery of islet function observed upon CR occurred while animals were maintained on HFD, albeit reduced caloric load. This prevented the deleterious effects of lipid overload on islet function and β-cell failure. Interestingly, studies show that CR can increase GLP1-Receptor (GLP-1R) expression in islets and favor the increase of PDX-1 expression, an important marker of β-cell maturation (Sheng et al., 2016). Exendin-4, a GLP-1R agonist, has been successfully used to improve glycemic control in patients with type 2 diabetes. Exendin-4 can also prevent the loss of Cx36 gap junction coupling in islets rendered dysfunctional by inflammatory cytokines (IL-1 and TNF-α) (Farnsworth et al, 2019). These findings reinforce that CR presents benefits to islet functionality by restoring the electrical coupling of β cells via Cx36 gap junctions.

In addition, in the present work we observed no changes in visibility if islet cells. This indicates that the results we observe for increases in islet dysfunction and functional recovery can be attributed to the effects of dietary modulation affecting islet function and not cell death. Thus CR can rapidly recover features associated with islet dysfunction during prediabetes, including Cx36 gap junction coupling and [Ca^2+^]_i_ dynamics.

In conclusion, we characterized the coordinated oscillations of [Ca^2+^]_i_, the permeability of Cx36 gap junction channels, and the overall insulin secretory function and glucose homeostasis of pre-diabetic mice subjected to CR. The decline in coordinated [Ca^2+^] and gap junction permeability at elevated glucose was diminished under high fat diet and fully rescued by short term CR. As such this shows how islet connectivity and secretory dynamics are diminished in pre-diabetes and can be rescued, thus contributing to the overall rescue of islet insulin secretion and glucose homeostasis.

## Acknowledgements

This study was partially supported by scholarship from the Brazilian funding agencies Fundação de Amparo à Pesquisa do Estado de São Paulo FAPESP, number 2018/04536-6 (to MACdA), as well as NIH grants R01 DK102950, R01 DK106412 and Juvenile Diabetes Research Foundation Grant 5-CDA-2014-198-A-N (to RKPB).

The authors would like to acknowledge the University of Colorado Anschutz Medical Campus Advanced Light Microscopy Core (supported in part by P30 NS048154, UL1 TR001082), as well as the Barbara Davis Center Islet Core for assistance with islet isolations.

